# Roux-Y Gastric Bypass and Sleeve Gastrectomy directly change gut microbiota composition independent of operation type

**DOI:** 10.1101/395657

**Authors:** Fernanda L. Paganelli, Misha Luyer, C. Marijn Hazelbag, Hae-Won Uh, Malbert R.C. Rogers, Danielle Adriaans, Roos-Marijn Berbers, Antoni P.A. Hendrickx, Marco C. Viveen, James A. Groot, Marc J. M. Bonten, Ad C. Fluit, Rob J.L. Willems, Helen L. Leavis

## Abstract

**Background:** Bariatric surgery in patients with morbid obesity, either through gastric sleeve gastrectomy or Roux-Y gastric bypass surgery, leads to sustainable weight loss, improvement of metabolic disorders and changes in the intestinal microbiota. Yet, the relationship between changes in gut microbiota, weight loss and the surgical procedure remains incompletely understood.

**Subjects/Methods:** We determined temporal changes in microbiota composition in 45 obese patients undergoing a crash diet followed by gastric sleeve gastrectomy (*n*= 22) or Roux-Y gastric bypass (*n*= 23). Intestinal microbiota composition was determined before intervention (baseline, S1), 2 weeks after a crash diet (S2), and 1 week (S3), 3 months (S4) and 6 months (S5) after surgery.

**Results:** Relative to S1, the microbial diversity index declined at S2 and S3 (*p*< 0.05), and gradually returned to baseline levels at S5. The crash diet was associated with an increased abundance of Rikenellaceae and decreased abundances of Ruminococcaceae and Streptococcaceae (*p*< 0.05). After surgery, at S3, the relative abundance of Bifidobacteriaceae had decreased (compared to the moment directly after the crash diet), whereas those of Streptococcaceae and Enterobacteriaceae had increased (*p*< 0.05). Increased weight loss during the next 6 months was not associated without major changes in microbiota composition. Significant differences between both surgical procedures were not observed at any of the time points.

**Conclusions:** In conclusion, undergoing a crash diet and bariatric surgery were associated with an immediate but temporary decline in the microbial diversity, with immediate and permanent changes in microbiota composition, with no differences between patients undergoing gastric sleeve gastrectomy or Roux-Y gastric bypass surgery.

## Introduction

Bariatric surgery is the only sustainable effective treatment for obesity ^1^. Surgical procedures such as the Roux-Y Gastric bypass (RYGB) and the sleeve gastrectomy (SG) facilitate a 50-70% decrease in excess body weight and fat mass ^1^. In addition, surgery leads to decreased caloric intake or malabsorption and to metabolic changes, such as an improved glucose metabolism, and is associated with a changed intestinal microbiota ^2-4^. The role of altered host-microbial interactions in this process is incompletely understood ^1^. Studies on the composition of the distal gut microbiota in obesity and after RYGB in humans and rodents yielded long lasting changes in types and ratio of enteric bacteria ^3, 5-8^. Furthermore, transfer of the gut microbiota from RYGB-treated mice to non-operated, germ-free mice resulted in weight loss and decreased fat mass in the recipient animals ^5^. These findings support a direct effect of the microbiota on weight and adiposity. Recently Liu *et al*. demonstrate using metagenomic shotgun sequencing that the abundance of glutamate-fermenting *Bacteriodes thetaiotaomincron* is decreased in obese Chinese individuals and glutamate levels are increased ^9^. Weight loss by SG partially reversed metabolic and microbial alterations, including reduced abundance of *B. thetaiotaomicron* and increased serum glutamate ^9^.

To further elucidate the results of the entire bariatric surgery procedure on the intestinal microbiota composition we investigated sequentially collected stool samples from 45 morbid obese patients undergoing either RYGB or SG at five different time points before and after surgery.

## Subjects and Methods

### Ethics Statement

The study protocol was in accordance with the regulations of the Ethics Committee of Catharina Hospital Eindhoven.

### Study Design

In this observational study patients with morbid obesity were recruited from the Catharina Hospital Eindhoven out-patient obesity clinic between September 2014 and November 2014. All 45 patients fulfilled the criteria for bariatric surgery and were screened before surgery for eligibility by a team including a surgeon, dietician and a psychologist. Two weeks before the planned surgery, patients were subjected to a crash diet consisting of 500 calories a day for 2 weeks. Type of surgery was determined based on clinical criteria and shared decision making between surgeon and patient. Reasons to refrain from RYGB were medication dependence, increased operation risk or super obesity. A reason to refrain from SG was gastroesophageal reflux disease. During surgery, patients received 1 g cefazolin antibiotic prophylaxis intravenously. After hospitalization, general practitioners managed adjustments of insulin, oral diabetics and other medication in the home setting. Patients visited out-patient clinic at 3, 6 and 12 months for evaluation and will remain in follow up for 5 years.

### Sample collection and DNA extraction

Stool samples (Sterilin specimen container, Thermo-Fisher) were gathered at the out-patient clinic or at patient homes. Samples were always stored in the freezer and collected at the homes of the patients using dry-ice and stored at the hospital at -80°C. Sample were collected at 5 different time points; before the start of the crash diet (S1), 2 weeks after the crash diet (S2), and 1 week (S3), 3 months (S4) and 5 to 6 months after surgery (S5).

Total bacterial DNA from feces samples was isolated according to Godon *et al*. ^10^. When isolated DNA contained PCR inhibitors (20% of the samples random distributed over the time points), samples were submitted to an extra step of isopropanol precipitation and column purification with QiAamp stool mini kit (Qiagen). DNA was stored at −20°C prior to further analysis.

### 16S rRNA gene sequencing strategy and analysis

A 469 bp encompassing the V3 and V4 hypervariable regions of the 16S rRNA gene was amplified and sequenced using the Illumina MiSeq Reagent Kit v3 (600-cycle) on an Illumina MiSeq instrument according to Fadrosh *et al*. ^11^. Negative controls, buffer controls were included in the DNA extraction, amplification and sequencing protocol to monitor for potential contamination. A total of 3 amplicon pools were sequenced, generating 8.9, 7.8 and 14.4 (mean of 10.3) million total reads. These 2x300 bp paired-end reads were pre-processed as follows. The first 12 bp of each paired-end containing the index sequences were extracted and afterwards concatenated to dual-index barcodes of 24 bp specific for each read-pair and sample. Paired reads were merged, as an overlap of about 90 bp was expected, using FLASH (version 1.2.11) ^12^. Subsequently, these merged reads were de-multiplexed using the split_libraries_fastq.py script rom and analyzed by the QIIME microbial community analysis pipeline (version 1.9.1) ^13^. Quality filtering was also performed during this step, truncating reads with an average PHRED quality score of 20 or less. After removal of the barcodes, heterogeneity spacers, and primer sequences about 19.8 million sequences were left with a mean length of 410 bp (median length of 405). The obtained sequences with a minimum of 97% similarity were assigned to operational taxonomic units (OTUs) using QIIME’s open-reference OTU picking workflow (pick_open_reference_otus.py). This workflow was carried out using USEARCH (version 6.1.544) ^14^ for OTU picking, in addition to detection and removal of chimeric sequences. The obtained OTU sequences were aligned to the Greengenes 16S rRNA gene database (gg_13_8_otus), followed by removal of OTUs represented by less than 0.005% of the total number of sequences. The generated OTU table and phylogenetic tree were used for assessing alpha- and beta-diversity using QIIME’s core_diversity_analyses.py workflow with a rarefaction depth of 20001 sequences. The weighted UniFrac distance was used to calculate beta-diversity of the samples, while the shannon index was used for the alpha-diversity. For Principal Component Analysis (PCA) R 3.5.0 in an environment of RStudio 1.1.383 (RStudio Team, Boston, MA) ^15^ was employed, using zCompositions, clr transformation and ggpplot R packages ^16-18^.

### Statistical analysis

Microbiota changes between time points and operation types were investigated using ANCOM ^19^ in R 3.3.3 ^15^. Changes in the clinical parameters (BMI, vitamin D, vitamin B6, cholesterol, bilirubin, glycated hemoglobin (HbA1c), iron, ferritin and folate) between baseline and 6 months after surgery was analyzed by applying t-test in Prism GraphPad (version 7.0). Associations between changes in total read counts at family level (at baseline versus 6 months after surgery) and changes in patient characteristics (at baseline versus 6 months after surgery) were investigated using a linear regression model. To eliminate possible confounding effects, age and sex were included as covariates. For these analyses, changes in total read counts were used as outcome, whereas changes in patient characteristics were used as predictor (model: change_in_read_counts ∼ β1·age + β2·sex + β3·change_in_patient_characteristic). For association analysis R 3.5.0 in an environment of RStudio 1.1.383 (RStudio Team, Boston, MA) was employed ^15^. Results are presented using pheatmap package (https://CRAN.R-project.org/package=pheatmap). For statistical testing we used false discovery rates (FDRs) correct for multiple comparisons, and an FDR- adjusted p-value < 0.05 was considered as significant ^20^.

## Results

### RYGB and SG resulted in significant decrease of BMI in all patients

In this study, 45 Caucasian Dutch patients were included with an average age of 43 years, 36 (84%) being female, 11 (24%) using proton pump inhibitors and 4 (9%) having type 2 Diabetes Mellitus at baseline (Table 1). After a crash diet, 22 patients underwent SG and 23 underwent RYGB. At baseline the mean BMI was 42.9 (+/-6.56) and 43 (+/-4.13) for patients undergoing RYGB and SG, respectively. At 6 months after the procedure BMI declined to 30.81 (+/-5.35) and 31.52 (+/-3.86), respectively, with no significant difference based on surgery type (Table 1).

### Crash diet reduces microbial alpha diversity, which is restored to baseline levels 6 months after surgery, irrespective of surgery type

In total 221 fecal samples were collected, with 4 samples missing from 4 unique times points from 4 different patients. Using a pre-defined cut-off value of 20001 reads, 220 samples could be analyzed.

The initial crash diet had a stronger effect on total microbiota diversity as the Shannon diversity index declined from 4.5 at baseline (S1) to 4.0 after the crash diet (*p*< 0.05) (S2) and then gradually returned to 4.5 at 3 (S4) and 6 months (S5) after surgery (Figure 1a). Differences in diversity are reflected by an initial decrease and subsequent rise in numbers of distinct microbial OTUs. At baseline, 3 months and 6 months after surgery more than 500 OTUs were identified, whereas after the crash diet and at 1 week after surgery this number was reduced to below 400 OTUs (Figure 1b).

In Principle Component Analysis (PCA) (Figure 1c), patients at baseline (S1) and after crash diet (S2) are more similar to each other when compared to the times points after the surgery (S3, S4, S5), which cluster together further apart.

**Figure 1.**
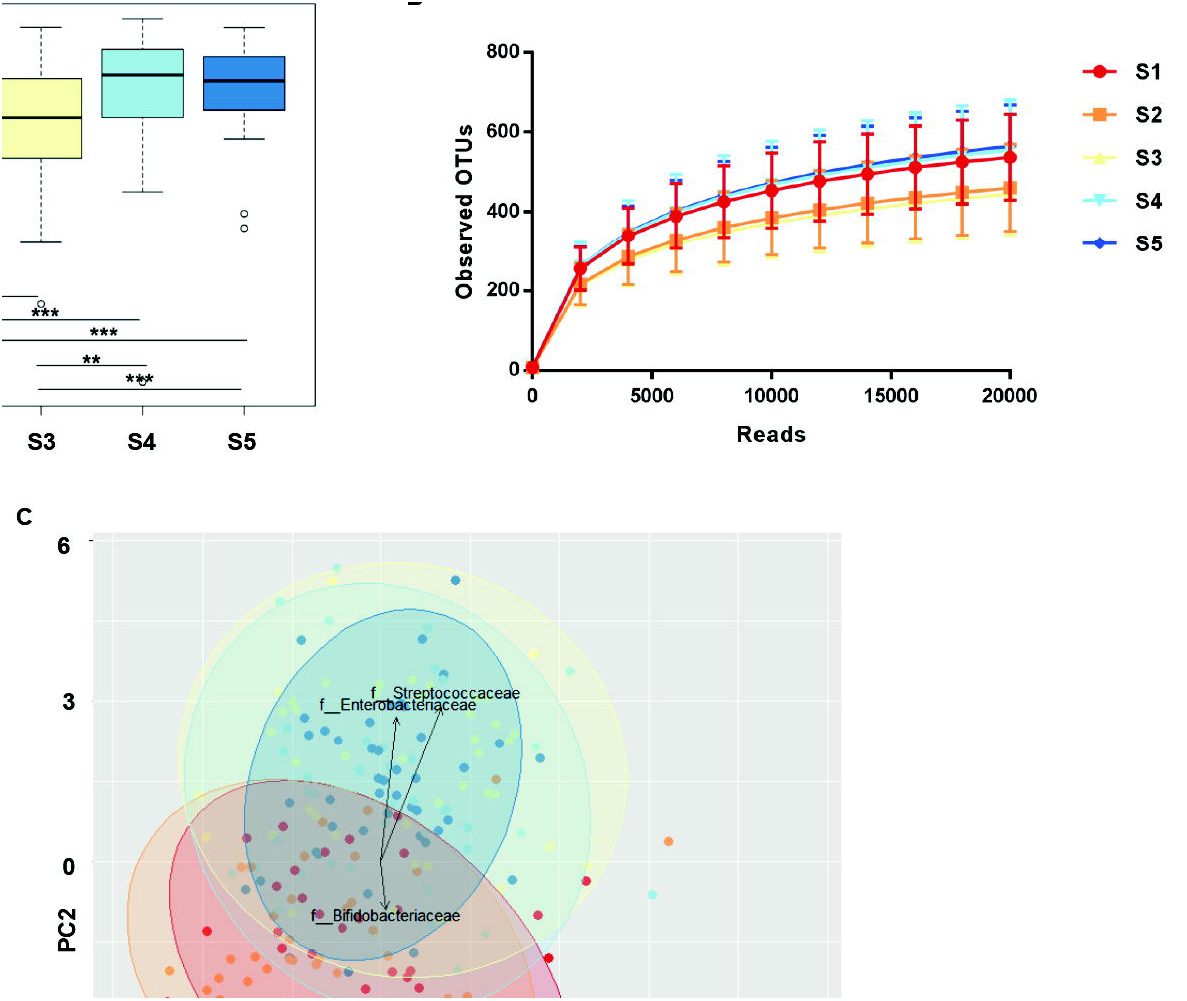
Observed and estimated richness of gut microbiota at different time points during the bariatric surgery procedure. (A). Shannon diversity index estimated a decrease in bacterial richness at S2 and S3. (B). Rarefication curves showed a reduction in bacterial richness at S2 and S3. (C). Principal component analysis (PCA) plot of similarity between the samples; each dot represents 1 sample, each color a different time point. S1. before surgery (red); S2. after 2 weeks of crash diet (orange); S3. 1 week after surgery (yellow); S4. 3 months after surgery (light blue); S5. 6 months after surgery (dark blue).

### Distinct microbial changes appear directly after crash diet, but are replaced by persistent distinct changes shortly after surgery

Significant changes in total relative abundance of specific families in the different time points were observed (Figure 2h). After the crash diet (S2) there was a significant reduction in relative abundance of 2 microbial families, Streptococcaceae and Ruminococcaceae (Figure 2a and d), and a significant increase in 1 family, Rikenellaceae (Figure 2e). Subsequent comparison of the microbial composition pre-surgery (S2) and 1-week post-surgery (S3) revealed a significant increase in the relative abundance of Streptococcaceae and Enterobacteriaceae families (Figure 2a and b) and a decrease in Bifidobacteriaceae, which persisted until 6 months post-surgery (S5) (Figure 2c). In these 6 months (at S5) microbiota complexity was restored (Figure 1a and b), which when compared with 1 week after surgery (S3) coincided with increased relative abundance of low abundance families Veillonellaceae and the Clostridiales order with no further family classification (Figure 2f and g).

**Figure 2.**
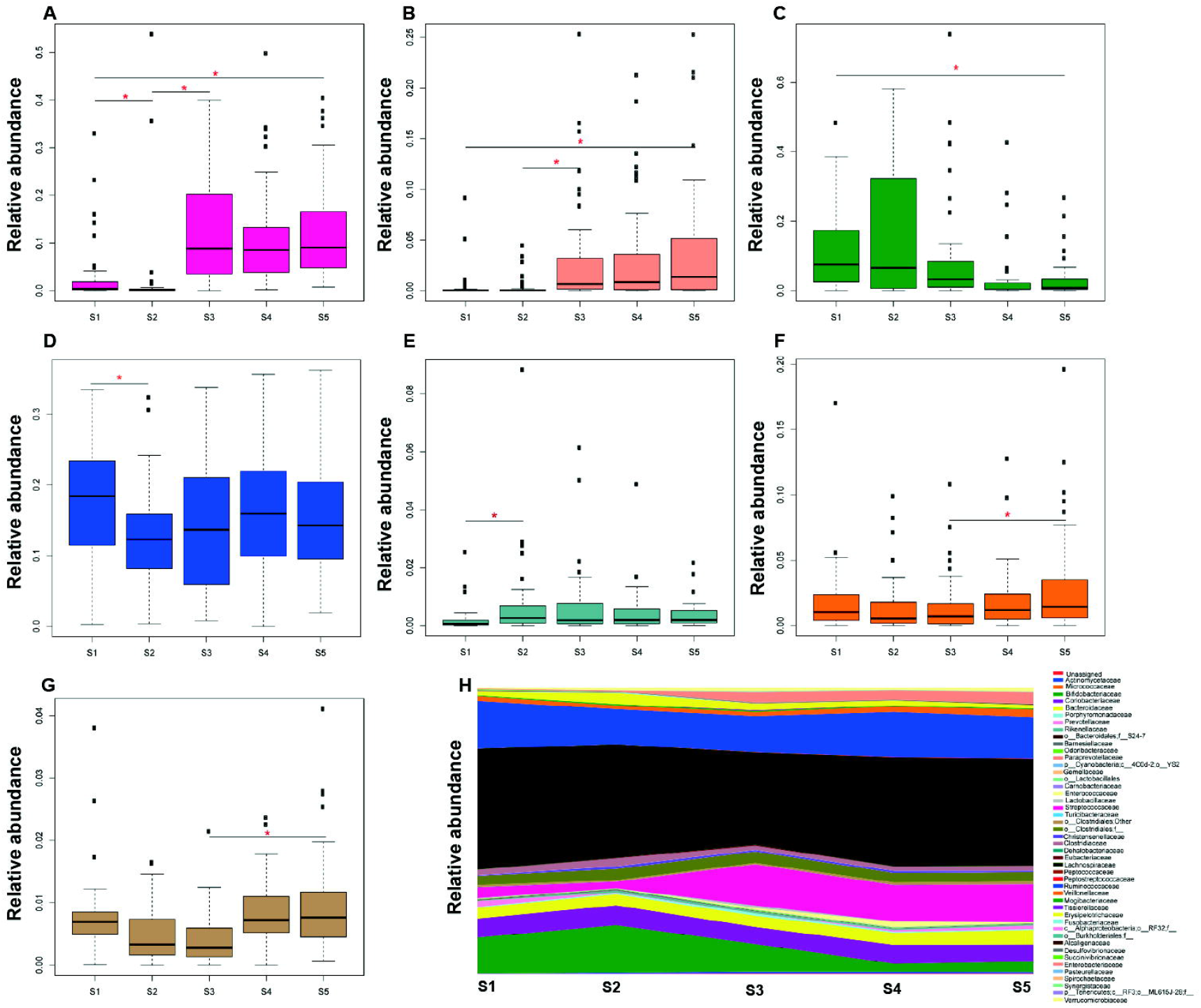
Relative abundance of bacterial families in the gut microbiota at the five time points analyzed. (A-G). Boxplots show the average relative abundance of 7 families that significantly changed between 2 different time points. (A). Streptococcaceae. (B). Enterobacteriaceae. (C). Bifidobacteriaceae. (D). Ruminococcaceae. (E). Rikenellaceae. (F). Veillonellaceae. (G). O_Clostridiales_f_others. H. Relative abundance of all families identified at the different time points. Significant families are represented in the same color. Asterisk (in red) indicates significant fold change differences (*p* < 0,05) analyzed by ANCOM.

When RYGB and SG surgeries were analyzed separated, no significant differences in microbiota composition based on beta diversity and relative abundance was observed at baseline (S1) (Figure 3a), 1 week (S3) (Figure 3b) or 6 months after surgery (S5) (Figure 3c) between patients that underwent either SG and RYGB (Figure 3d).

**Figure 3.**
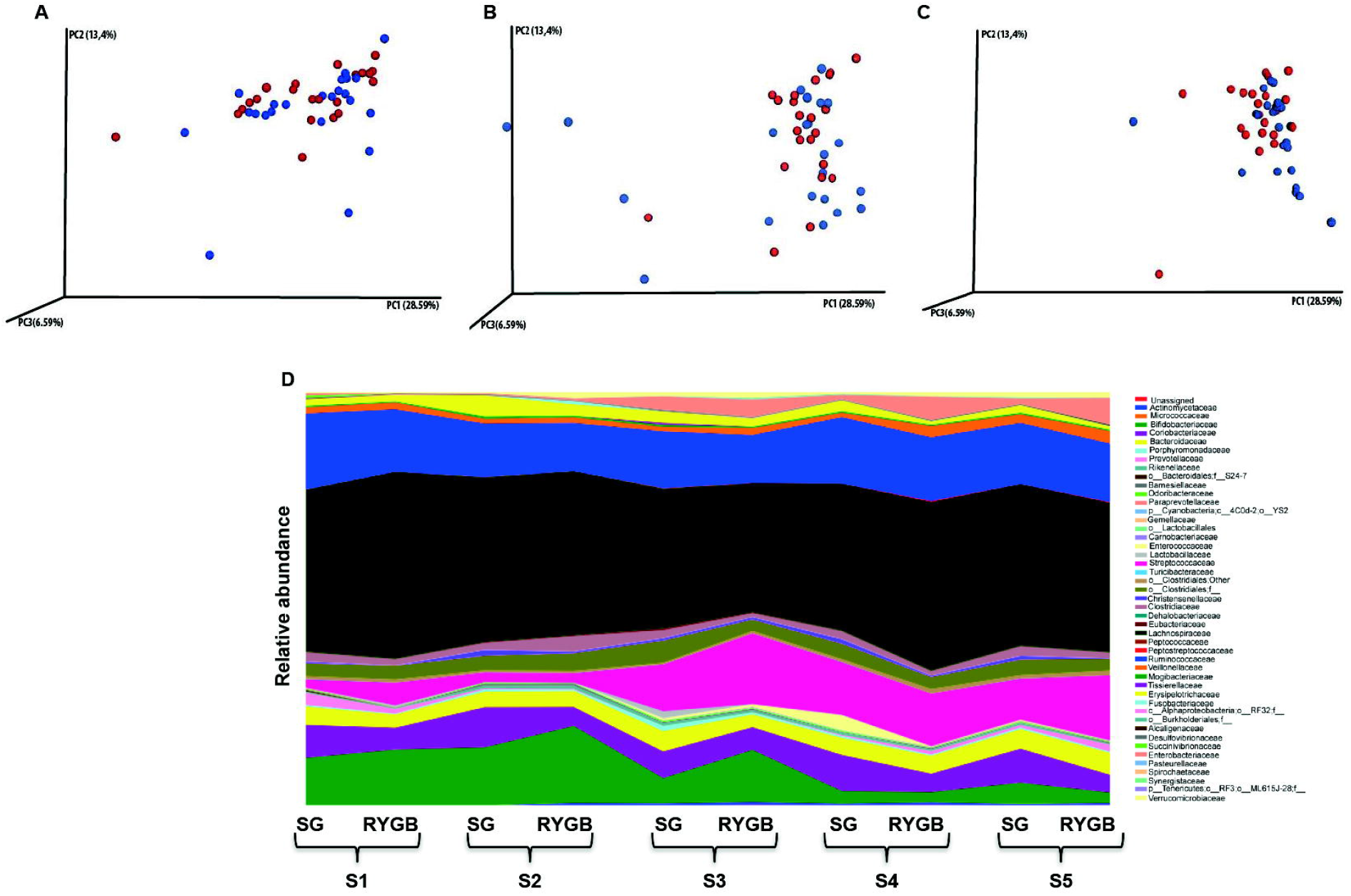
(A-C). Principal coordinate analysis (PCoA) plots comparing beta diversity of Sleeve Gastrectomy (SG) versus Roux-Y Gastric bypass (RYGB) surgery at baseline (S1) (A) 1 week after surgery (S3) (B) and 6 months after surgery (S5) (C). SG is indicated in red, RYGB is indicated in blue (D). Relative abundance of bacterial families in the gut microbiota at the five time points analyzed in SG versus RYGB surgery.

### Significant associations between microbiota changes and clinical markers

Clinical parameters in patients were analyzed at baseline and 6 months after the surgery. Besides weight loss, serum levels of vitamin D, B6, cholesterol, bilirubin, HbA1c, iron, ferritin and folate improved 6 months after surgery when compared to baseline (FDR-adjusted, *p*< 0.05, Table 1).

These changes were associated to overall differences in microbial abundance in relation to the changes in clinical parameters at S5 versus S1, which are highlighted in Figure 4. Significant associations were only found in low abundance families. Decreased bilirubin level was associated with and Prevotellaceae, Bacteroidales and Peptococcaceae taxa; and increased iron level was associated with Pasteurellaceae. In addition, a decreased HbA1c was associated with Coriobacteriaceae and Clostridiales taxa The most pronounced measured effects in the dataset was a negative association between Prevotellaceae, Veillonellaceae, Streptococcaceae, Bifidobacteriaceae and Enterobacteriaceae taxa in relation to decreased serum cholesterol levels, whereas most pronounced positive associations were found between Lachnospiraceae and Coriobacteriaceae taxa in relation to decreased cholesterol levels (Figure 4). Yet, these associations were not statistically significant after FDR adjustment (Figure 4).

**Figure 4.**
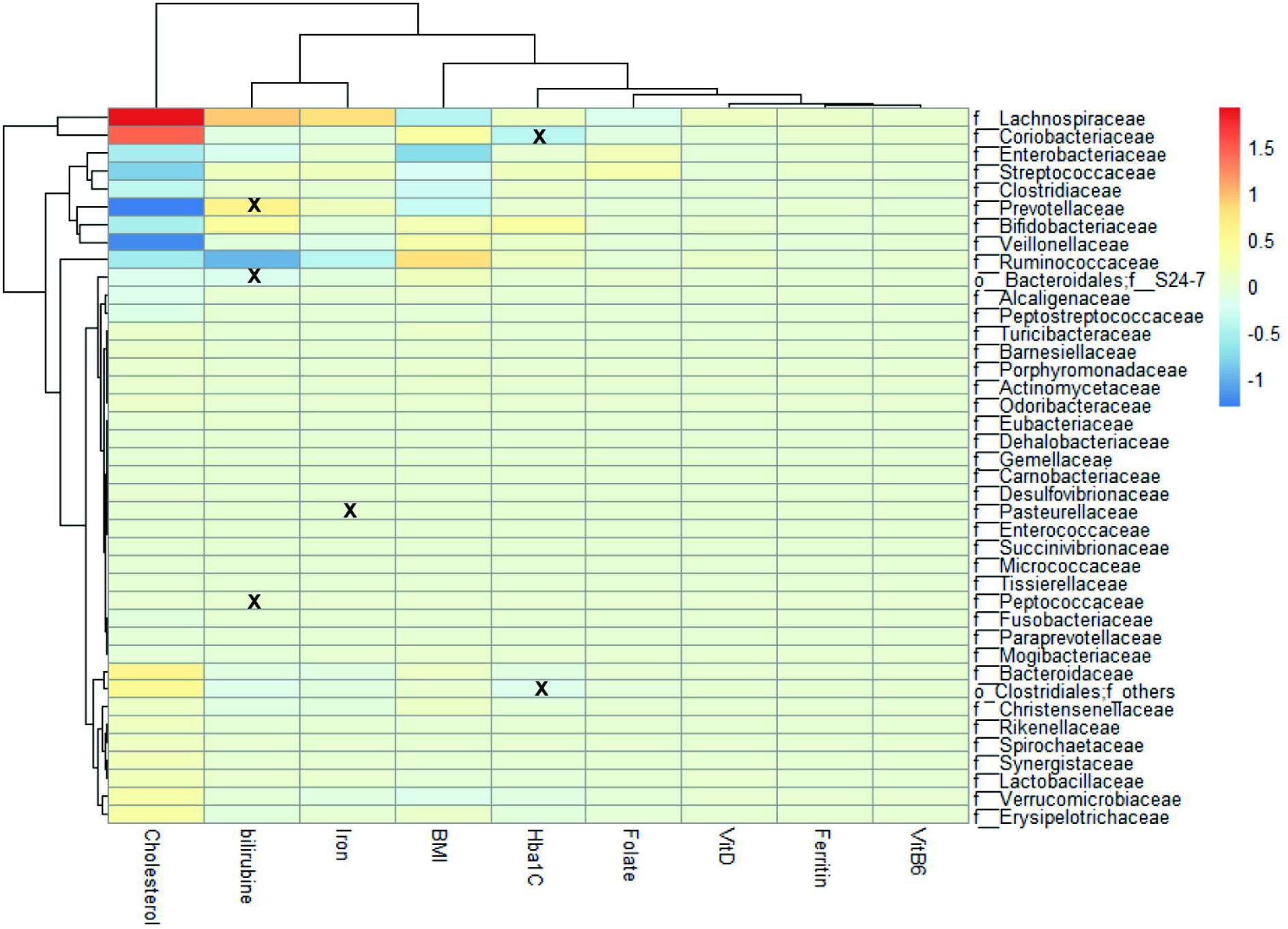
Association between clinical parameters and family taxa calculated based on the difference between 6 months after surgery (S6) and baseline (S1). The significant association (false discovery rate (FDR) adjusted p-value<0.05) is indicated with a “x”. The red color indicates positive effect and the blue color negative effect. HbA1c, glycated hemoglobin; VitD, vitamin D; VitB6, vitamin B6.

## Discussion

This study is novel in the fact that it compares microbiota profiles before and very shortly after bariatric surgery with subsequent follow-up profiles. We describe the sequential impact of a crash diet followed by either RYGB or SG surgery, resulting in progressive weight loss and changes in the gut microbiota composition. Several other studies report sequential sampling of patients after bariatric surgery, but none of the studies define timing of the baseline sample in relation to a crash diet ^3, 9, 16, 21-23^, and, therefore, renders the relative impact of the different measures in the bariatric procedure difficult to dissect. In addition, this study is unique in that a very early postsurgical sampling time point is included. Apart from substantial weight loss and improvements in clinical parameters, as reported by others, bariatric surgery induces long-lasting changes in microbiota composition in most patients. The most apparent immediate change in microbiota composition occurred after the crash diet, with a concurrent reduction in alpha diversity, whereas surgery was associated with early and sustained replacement of distinct bacterial taxa and restoration of the diversity.

Although significant microbial changes are identified in the gut microbiota of bariatric surgery patients, at 6 months after surgery the total microbial diversity was similar to microbial diversity measured at baseline. This sudden decline in alpha diversity probably reflects a severe stress on the human microbiota by a crash diet, with a significant change in catabolic state ^24^. Persisting post-surgery microbiota changes suggest adaptation to anatomic and physiologic changes, such as reduced acid production, increased oxygen content, altered bile acid concentrations delivered to the colon, induced by the surgery. Previously reported effects of bariatric surgery on microbiota diversity have ranged from an increase in total diversity ^25, 26^ to absence of change and even a decrease in alpha diversity ^27^. We suspect that baseline sampling in relation to crash diet may vary between studies, and might contribute to the reported differences between studies.

Besides a stable bacterial alpha diversity after surgery, we observed profound differences after each consecutive intervention on bacterial taxa composition. The crash diet immediately resulted in an increase in the relative abundance of Bifidobacteriaceae and decrease of Streptococcaceae, whereas the opposite effect was observed after surgery; an increase in abundance of Streptococcaceae and decline in Bifidobacteriaceae that persisted for at least 6 months. In addition to the observed increased abundance of Veillonellaceae may reflect survival of oral microbiota into the intestine. Observed persistent increase in Enterobacteriaceae after surgery, confirms previous sustained changes reported in humans and animal models (rats), associated with increased pH ^7, 24, 28, 29^. Other main differences, exposure to undigested nutrients and biliopancreatic enzymes, may play important roles in the microbial composition, intestinal permeability and intestinal adaptation ^30^. Since increased intestinal permeability is associated with inflammation and reductions in alpha diversity, which is also associated with obesity, it is questionable whether restoration of alpha diversity to baseline level may also reflect persistent inflammation in the post-surgery state at 6 months, which has been previous related to increase in Enterobacteriaceae ^16^. This corresponds to the observed higher alpha diversity of fecal samples from a healthy normal weight cohort compared to the although improved, yet lower diversity of postoperative patients ^16^.

Although others observed microbiota changes only after RYGB ^5^, here we observed this in both surgery types. This suggests that despite the 2 procedures result in distinct anatomic differences, this did not seem to influence the post-surgery changes in relative abundance of Bifidobacteriaceae, Streptococcaceae and Enterobacteriaceae observed amongst both patient groups and which were similar for both types of surgery. Interestingly, unlike Liu *et al*. ^9^ and Ilhan *et al*. ^31^ both patients groups here after surgery develop comparable weight loss irrespective of surgery type, this may explain why we find similar changes in gut microbiota composition. Also baseline characteristics did not differ significantly. Moreover, we suggest that bariatric surgery in itself, unlike crash diet, results in an altered long-lasting composition of the microbiota.

Although a significant association with changed clinical parameters between baseline and 6 months after surgery was lacking, the relative abundance of Bifidobacteriaceae, Streptococcaceae and Enterobacteriaceae taxa changed significantly shortly after surgery. This sudden adjustment further confirms that the altered postoperative microbiota more likely reflects surgery induced effects, rather than improved clinical parameters ^16, 23, 31^. We observed a significant association between increased serum bilirubin level and decreased relative abundance of Bacteroidales, Peptococcaceae and Prevotellaceae taxa in this dataset. The abundance of Bacteroidales in the gut microbiota could contribute to the increase in bilirubin level, since *Bacteroides fragilis*, which is part of Bacteroidales taxa, is one of the bacterial species described to be able to metabolize bilirubin in the gut ^32, 33^. In addition, a decreased HbA1c was found significantly associated with decreased Coriobacteriaceae and increased Clostridiales taxa. Nevertheless, the exact meaning of changes of these low abundance taxa is unknown.

This study failed to confirm the suggested relationship between increased abundance of Firmicutes and Bacteroidetes and obesity ^34, 35^, as the relative abundance of the family members of these phyla remained stable before and after surgery, despite significant weight loss. In addition, other studies described that Faecalibacterium (*F. prausnitzii*) was assumed to play a role in inflammation and glucose homeostasis in obesity with a reduced relative abundance after RYGB surgery ^3, 8, 36, 37^. In our study, a decreased abundance of the Ruminococcaceae family, to which *F. prausnitzii* belongs, was observed after the crash diet, yet this change did not sustain after surgery.

In conclusion, here we illustrate that temporal sampling of bariatric surgery patients with subsequent microbiota analysis can lead to increased insights into the relative contribution of interventions on stability and composition of the microbiota. We show that a crash diet invoked profound temporary changes in total microbiota diversity and composition, yet surgery precluded early fixed changes of microbial composition and restoration of the microbial diversity that likely contribute to weight loss.

## Acknowledgments

The authors declare that there are no conflicts of interest. Authors contribution: conceived and designed the experiments: ML, DA, MJMB, ACF, HLL. Performed the experiments: ML, DA, APAH, JG, MCV. Analyzed the data: FLP, MRCR, CMH, HWU, RMB. Wrote the paper: FLP, ML, CMH, HWU, MRCR, DA, RMB, APAH, MCV, JG, MJMB, ACF, RJLW, HLL.

## Competing interests

The authors declare no competing financial interests.

